# Virus-induced gene silencing (VIGS) as a tool for functional genetic analysis in passion fruit plants

**DOI:** 10.1101/2023.11.27.568936

**Authors:** Xiaoqing Wang, Wentao Shen, Hongguang Cui, Zhaoji Dai

**Affiliations:** Sanya Nanfan Research Institute, Key Laboratory of Green Prevention and Control of Tropical Plant Diseases and Pests (Ministry of Education), School of Plant Protection, Hainan University, Haikou, Hainan, China; Collaborative Innovation Center of Nanfan and High-Efficiency Tropical Agriculture, Hainan University, Haikou, Hainan, China; Hainan Key Laboratory for Protection and Utilization of Tropical Bioresources, Institute of Tropical Bioscience and Biotechnology, Sanya Research Institute, Hainan Institute for Tropical Agricultural Resources, Chinese Academy of Tropical Agricultural Sciences, Haikou & Sanya, Hainan, China

**Keywords:** Passion fruit, *Passiflora edulis*, Passiflora, Virus-induced gene silencing, VIGS, telsoma mosaic virus, TelMV, *phytoene desaturase*, PDS

## Abstract

Passion fruit (*Passiflora edulis*) is grown perennially in sub-tropical and tropical areas. Its fruits contain multiple vitamins and antioxidants, and thus are consumed increasingly in drinks and foods. However, the functions and regulations of genes that are engaged in the biosynthesis of the health-promoting compounds in passion fruits remain largely unknown. Its whole genome sequence has just been published recently. Virus-induced gene silencing (VIGS) is a reverse genetics tool for analyzing gene function. Here, we engineered telosma mosaic virus (TelMV), a potyvirus infecting passion fruit, into a VIGS vector by inserting the Gateway-compatible recombination sites. The newly constructed TelMV-VIGS virus successfully expressed foreign protein and induced systemic infection in both *Nicotiana benthamiana* and *P. edulis* plants. Intriguingly, TelMV-VIGS vector containing different fragments of green fluorescent protein (GFP) gene induced systemic gene silencing on the GFP-transgenic *N. benthamiana* plants (16c). When the *phytoene desaturase* (*PDS*) gene, an endogenous gene in passion fruit, was engineered into the vector, it triggered the silencing of *PePDS*, as evidenced by the reduced mRNA levels and photobleached phenotype. We reported the first development of VIGS vector in passion fruit, as the first step in our endeavor of discovering horticulturally important genes for improving passion fruit production and quality.

## Introduction

Plant viruses can be used as vectors for protein expression and virus-induced gene silencing (VIGS) as they comprise relatively small genomes, are easy to manipulate, and can spread throughout the whole plant (Constantin et al., 2004; Cui et al., 2017). As a reverse genetic tool, VIGS is a powerful alternative approach for determining gene function in species that are not amenable to the classic genetic transformation methods, such as *Agrobacterium*-mediated transformation (Constantin et al., 2004; Hileman et al., 2005). In addition, VIGS possess the great advantage of being fast that the time from cloning the gene of interest (GOI) to the phenotype analysis is much shorter compared to stable plant transformation. Typically, VIGS involves the cloning of fragments of GOI into the viral vector, viral infection of the plant hosts, and silencing of the target genes by the RNA-mediated antiviral defense mechanism in plants (Rossner et al., 2022). Since the first report of VIGS in 1995 (Kumagai et al., 1995), VIGS has been proven to be an efficient tool for gene function analysis in diverse plant species, including tobacco, soybean, wheat, tomato, Arabidopsis and petunia (Dommes et al., 2019; Rossner et al., 2022).

Passion fruit (*Passiflora edulis*) is a perennial evergreen vine grown in sub-tropical and tropical areas. It belongs to the *Passiflora* genera in the family of *Passifloraceae* (Xia et al., 2021; Wang et al., 2023). Over the years passion fruits have become the new favorites in both drinks and foods as it contains multiple vitamins and antioxidants that benefit human health. However, the functions and regulations of genes that are engaged in the biosynthesis of the health-promoting compounds in passion fruits remain largely unknown. This is mainly due to the lack of efficient tools for gene function analysis. In 2021, a chromosome-scale genome assembly of passion fruit (*Passiflora edulis* Sims) was just reported (Xia et al., 2021) and the gene function study of passion fruit is very much in its infancy. Despite the fact that the in vitro regeneration and the *Agrobacterium*-mediated stable transformation of passion fruit have been established (Trevisan et al., 2006; Correa et al., 2015; Rizwan et al., 2021), the process are time-consuming and inconvenient for high-throughput characterization of passion fruit genes. An alternative approach is the use of Gateway technology-based VIGS. In this system, the Gateway recombination cassette *att*R1-Cm^R^-*ccd*B-*att*R2 is inserted in the viral vector and thus allows the simple, fast and high-throughput cloning of the gene of interest. Multiple viral vectors compatible with the Gateway cloning technology have been reported for protein expression and the inducing of VIGS in plants, including tobacco rattle virus (TRV), potato virus X (PVX), tobacco mosaic virus (TMV), and cymbidium mosaic virus (CymMV) (Liu et al., 2002; Lacorte et al., 2010; Lu et al., 2012). The most famous and broadly-used VIGS vector is the TRV-based vector. In 2002, Liu et al. reported the Gateway-compatible TRV VIGS vector that is successfully applied to silence endogenous gene in tomato (Liu et al., 2002).

Telosma mosaic virus (TelMV) is an emerging plant virus infecting passion fruit plants in multiple plantations in China, Brazil, Vietnam and Japan (Gou et al., JVI, 2023). TelMV belongs to the genus *Potyvirus* in the family *Potyviridae*. Potyvirus represents the largest group of plant RNA viruses and comprises a positive-sense single-stranded RNA genome encapsidated by viral coat protein. The potyvirus genome encodes a large polyprotein that is further cleaved into ten mature viral proteins by its own viral proteases (Revers and García, 2015; Cui and Wang, 2019; Dai et al., 2020; Yang et al., 2021). Although potyviruses have been broadly engineered into protein expression vectors, such as tobacco etch virus (TEV), potato virus A (PVA), turnip mosaic virus (TuMV), plum pox virus (PPV), sugarcane mosaic virus (SCMV), wheat streak mosaic virus (WSMV) and zucchini yellow mosaic virus (ZYMV) (Xie et al., 2021), very few potyvirus-based VIGS vector have been reported. The most common insertion site positions of these potyvirus-based vectors lie in the P1/HCPro and NIb/CP junctions (Xie et al., 2021; Houhou et al., 2021). Luckily, our lab has previously reported the first passion fruit-infecting TelMV infectious clone pPasfru (Gou et al., 2023).

Currently, the VIGS tool has not yet been reported in passion fruit. Thus, this study aims to develop a VIGS system in passion fruit plants using the potyvirus TelMV as a silencing vector. Here, we engineered a Gateway-compatible TelMV VIGS vector (TelMV-GW) for assessing the biological function of the endogenous gene in passion fruit. The TelMV-GW was revealed to systemically infect both *N. benthamiana* and passion fruit plants and expressed foreign protein upon insertion of foreign gene. Furthermore, TelMV-GW is capable of silencing the *P. edulis phytoene desaturase* (*PePDS*) gene, showing a photobleached phenotype in VIGS-treated passion fruit plants. The TelMV-derived VIGS tool described here constitutes an ideal tool for functional genetic analysis in passion fruit plant, especially the genes engaged in the biosynthesis of the health-promoting compounds in passion fruits.

## Results

### Development of telosma mosaic virus (TelMV)-based VIGS vector compatible with Gateway cloning technology

To develop a VIGS vector for passion fruit plants, we decided to engineer telosma mosaic virus (TelMV), a potyvirus infecting passion fruit, into a viral vector that is compatible with the conventional Gateway cloning technology. To this end, the Gateway recombination frame comprising the NIa-Pro cleavage sites and the *att*R1-*Cm*^*R*^*-ccd*B -*att*R2 recombination cassette was cloned by the overlapping PCR method as described in the Materials and Methods, and subsequently inserted between the NIb and CP cistrons in the backbone of TelMV infectious clone pPasFru (Gou et al., 2023) (Figure 1). The resulting construct was designated TelMV-GW and further verified by double digestion and Sanger sequencing. To determine whether TelMV-GW could systemically infect plants and express protein upon insertion of foreign gene, an entry clone containing the GFP gene was used for LR recombination reaction. The resulting construct was named TelMV-GW-GFP. Upon inoculation of *Agrobacterium* harboring TelMV-GW-GFP in the *N. benthamiana* plants, clear green fluorescence could be visualized in the upper non-inoculated leaves under UV illumination at 15 days post inoculation (dpi) (Figure 2A), same as the plants inoculated with the positive control, the previously reported GFP-tagged TelMV infectious clone (TelMV-GFP) (Gou et al., 2023).

**Fig 1.**
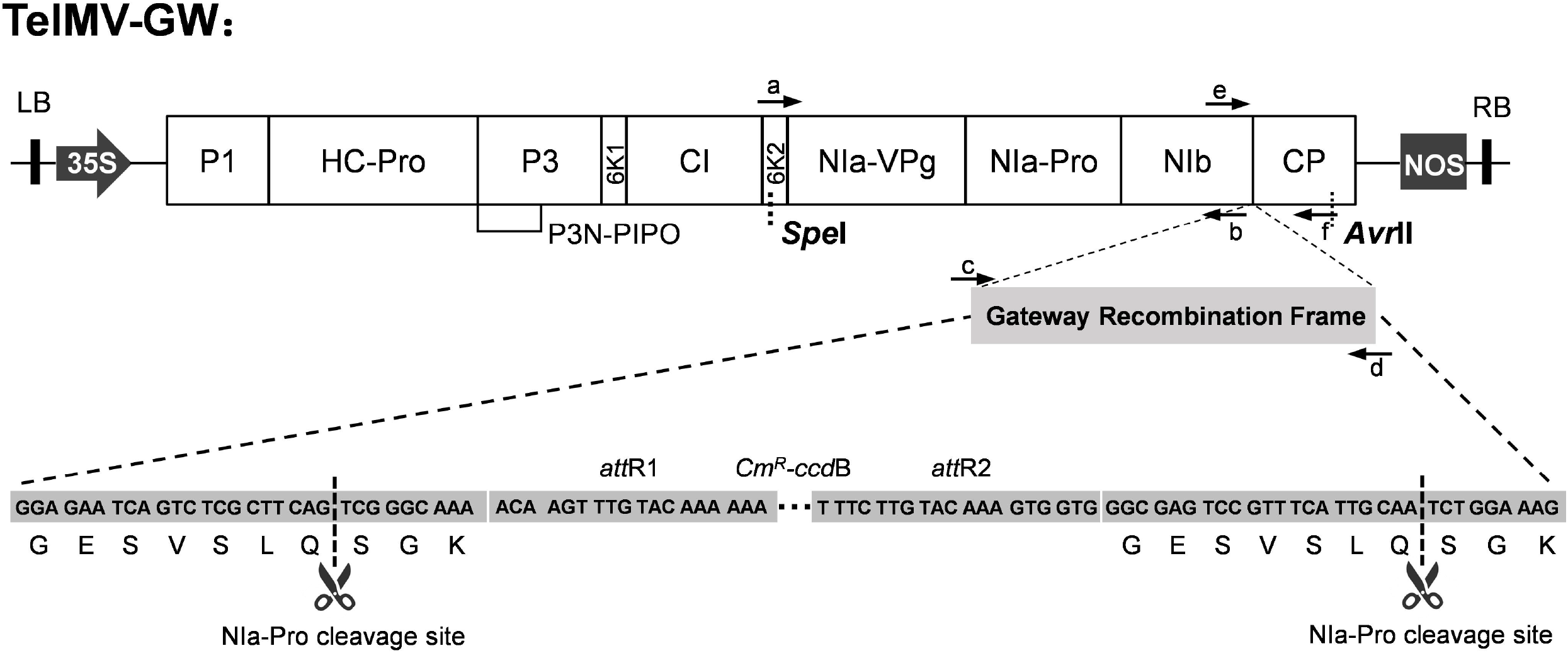
Schematic representation of the Gateway-compatible TelMV vector (TelMV-GW). The five letters (a, b, c, d, e, and f) represent the primers used for TelMV-GW cloning by overlapping PCR. The Gateway recombination frame was inserted in between the viral nuclear inclusion protein b (NIb) and coat protein (CP) cistrons. The magnification shows the Gateway recombination frame sequence that contains the last seven amino acid resides of NIb cistron (GESVSLQ), the first three amino acid resides of CP cistron (SGK) and the Gateway recombination cassette *att*R1-Cm^R^-*ccd*B-*att*R2. LB, RB: left and right border of T-DNA; 35S: 35S promoter of cauliflower mosaic virus; Nos: nopaline synthase terminator; *att*R1, *att*R2: Gateway recombination sites; Cm^R^, *ccd*B, selection markers for Gateway cloning.

**Fig 2.**
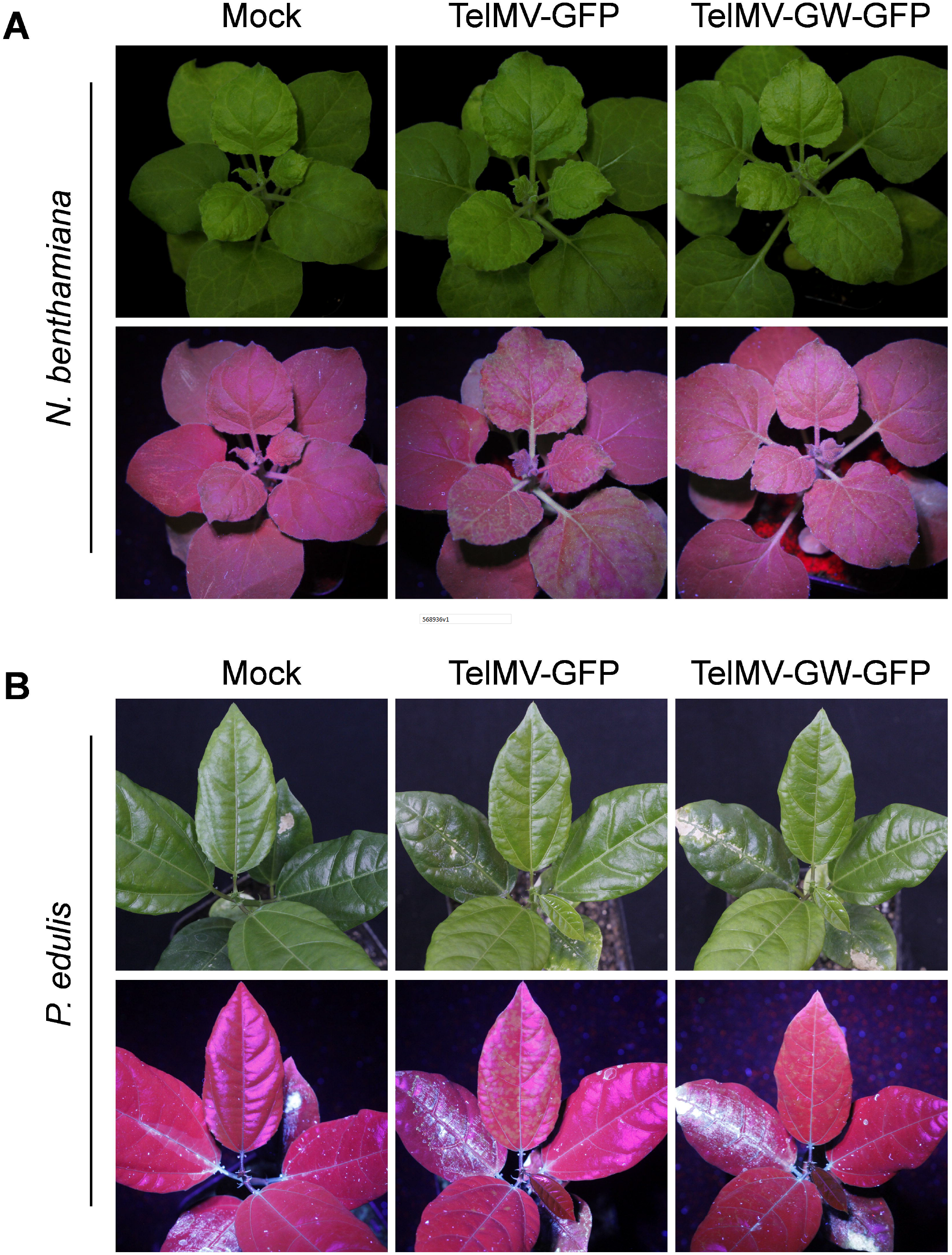
TelMV-GW as an expression vector in *N. benthamiana* and *P. edulis* plants. (A) *N. benthamiana* plants inoculated by buffer (Mock), positive control (TelMV-GFP) and TelMV-GW-GFP, respectively under UV light at 15 dpi. (B) *P. edulis* plants inoculated by buffer (Mock), positive control (TelMV-GFP) and TelMV-GW-GFP, respectively under UV light at 15 dpr. Note that clear green fluorescence could be observed on the newly emerged leaf of both *N. benthamiana and P. edulis* inoculated with TelMV-GW-GFP upon UV illumination.

More importantly, upon rub-inoculation of passion fruit plants with the sap prepared from TelMV-GW-GFP-infected *N. benthamiana* leaf at 15 days post rub-inoculation (dpr), intense green fluorescence was observed under UV light, suggesting TelMV-GW-GFP can systemically infect the passion fruit plants (Figure 2B). Furthermore, RT-PCR and Western blot analysis confirmed the presence of mRNA and protein of GFP (data not shown), respectively in the upper leaf of both *N. benthamiana* and passion fruit plants inoculated by TelMV-GW-GFP. These results demonstrated that TelMV-GW serves as an excellent expression vector in both *N. benthamiana* and passion fruit plants.

### Visualization of VIGS in the 16c *N. benthamiana* plants

To examine the potentiality of the Gateway-compatible TelMV vector (TelMV-GW) for virus-induced gene silencing (VIGS), the green fluorescent protein (GFP)-transgenic *N. benthamiana* plants (16c) were used for gene silencing analysis. The full length of GFP was divided into two parts, 309 bp and 405 bp, and subsequently cloned into the TelMV-GW vector. The resulting constructs were named TelMV-GW-GFP309 and TelMV-GW-GFP405, respectively. The 16c plants were agroinoculated with either TelMV-GW-GFP309, TelMV-GW-GFP405 or wild-type TelMV (TelMV). As expected, at 18 dpi, the 16c plants inoculated with TelMV-GW constructs carrying GFP fragments showed apparent red fluorescence with reduced green fluorescence in the upper leaves compared to the Mock- or TelMV-infiltrated plants (Figure 3A), suggesting the GFP gene is likely silenced in the TelMV-GW-GFP309 and 405-inoculated 16c plants. This phenomenon is much more obvious at 26 dpi where the systemic leaves of 16c plants inoculated by TelMV-GW constructs carrying GFP fragments exhibited mainly red fluorescence under UV light (Figure 3A), indicating the GFP gene is silenced to a great extent.

**Fig 3.**
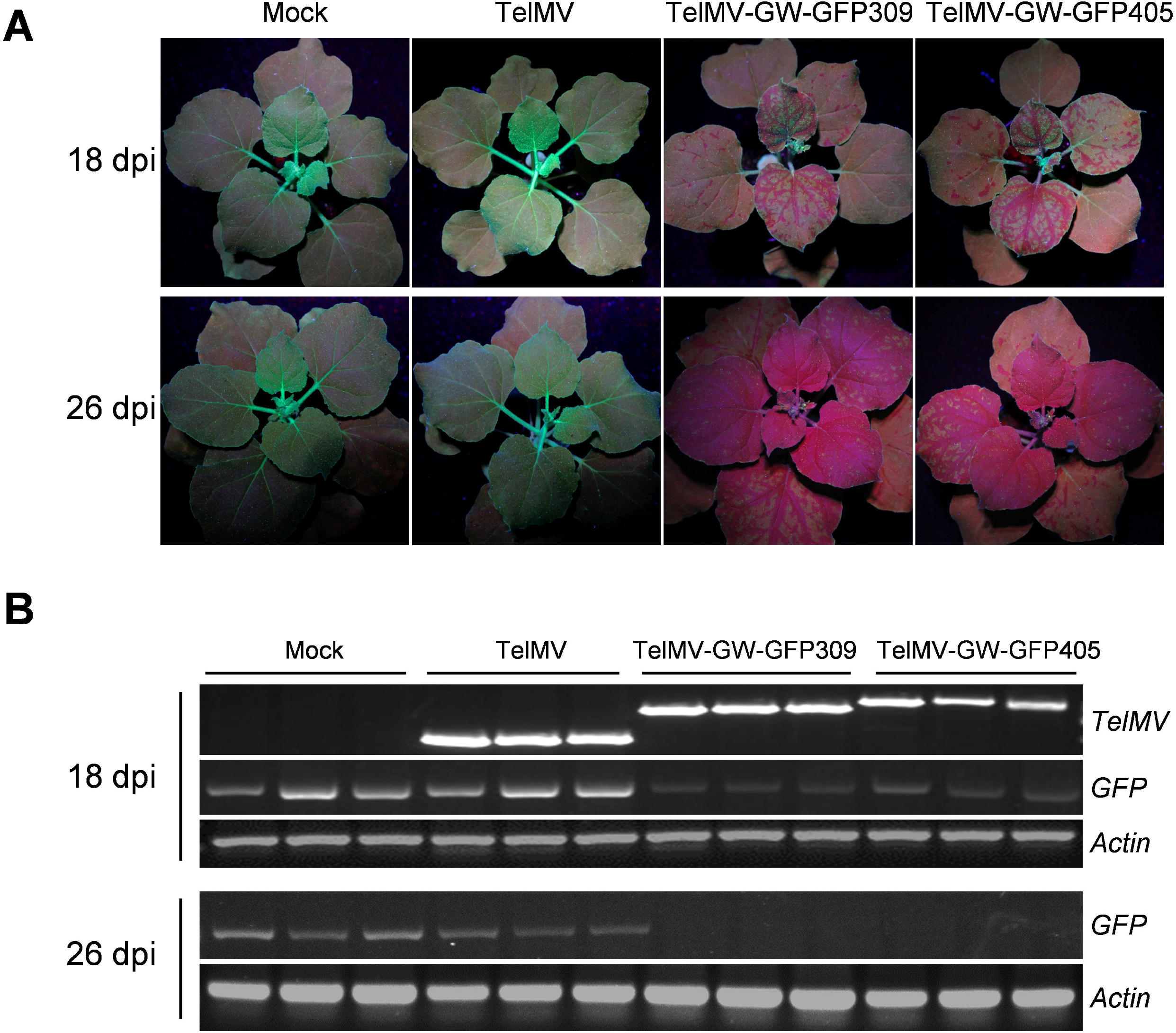
TelMV-based VIGS vector can trigger gene silencing in the GFP-transgenic *N. benthamiana* plants (16c). (A). 16c plants inoculated by the TelMV-GW constructs harboring different GFP fragments, Wild-type TelMV (TelMV) or buffer (Mock), under UV light at 18 and 26 dpi, respectively showing the progression of GFP-silencing induced by TelMV-GW VIGS vector. (B). RT-PCR analysis of the GFP mRNA level of 16c plants inoculated by various constructs at 18 and 26 dpi, respectively. TelMV-specific primers were used to detect viral RNA. The *Actin* gene serves as an internal control.

Next, we extracted total RNA from the upper leaf of each sample and performed RT-PCR analysis. The results revealed a reduced GFP mRNA level of *N. benthamiana* leaf inoculated by TelMV-GW-GFP309 or TelMV-GW-GFP405, compared to that of Mock- or TelMV-infiltrated plants at 18 dpi (Figure 3B). In addition, the GFP fragments are genetically stable in the systemic leaf evidenced by the amplicons of correct size using the primer pairs that can amplify bigger fragments comprising full-length GFP (Figure 3B). At 26 dpi, RT-PCR analysis showed a nearly undetectable mRNA level of GFP (Figure 3B), indicating the GFP gene of 16c plants was silenced upon inoculation of TelMV-GW constructs carrying GFP fragments. Taken together, we demonstrated that the TelMV-GW vector is capable of triggering gene silencing in the 16c plants.

### Silencing of *phytoene desaturase* (*PDS*) in passion fruit plants

The *phytoene desaturase* (*PDS*) gene is essential for carotenoid production and its silencing results in photobleaching. Thus, we chose *PDS* gene for the test of TelMV-induced endogenous gene silencing in passion fruit plants. A 345 bp fragment of the *PePDS, PePDS*_*345*_, was amplified from the cDNA of passion fruit leaf and cloned into the TelMV-GW, resulting in TelMV-GW-PDS_345_. Passion fruit plants in the first- to second-true-leaf stage were rub-inoculated with sap prepared from TelMV-GW-PDS_345_-inoculated *N. benthamiana* leaf. At 50 days post rub-inoculation (dpr), passion fruit plants inoculated with TelMV-GW-PDS_345_ showed photobleaching on the upper, uninoculated leaves (Figure 4A), indicating the endogenous gene *PDS* had been successfully silenced. By 90 dpr, more apparent photobleaching phenomenon can be observed in the systemic leaves of passion fruit plants inoculated with TelMV-GW-PDS_345_. In contrast, all leaves of plants inoculated with buffer (Mock) or TelMV-GW-GFP remained green (Figure 4A).

**Fig 4.**
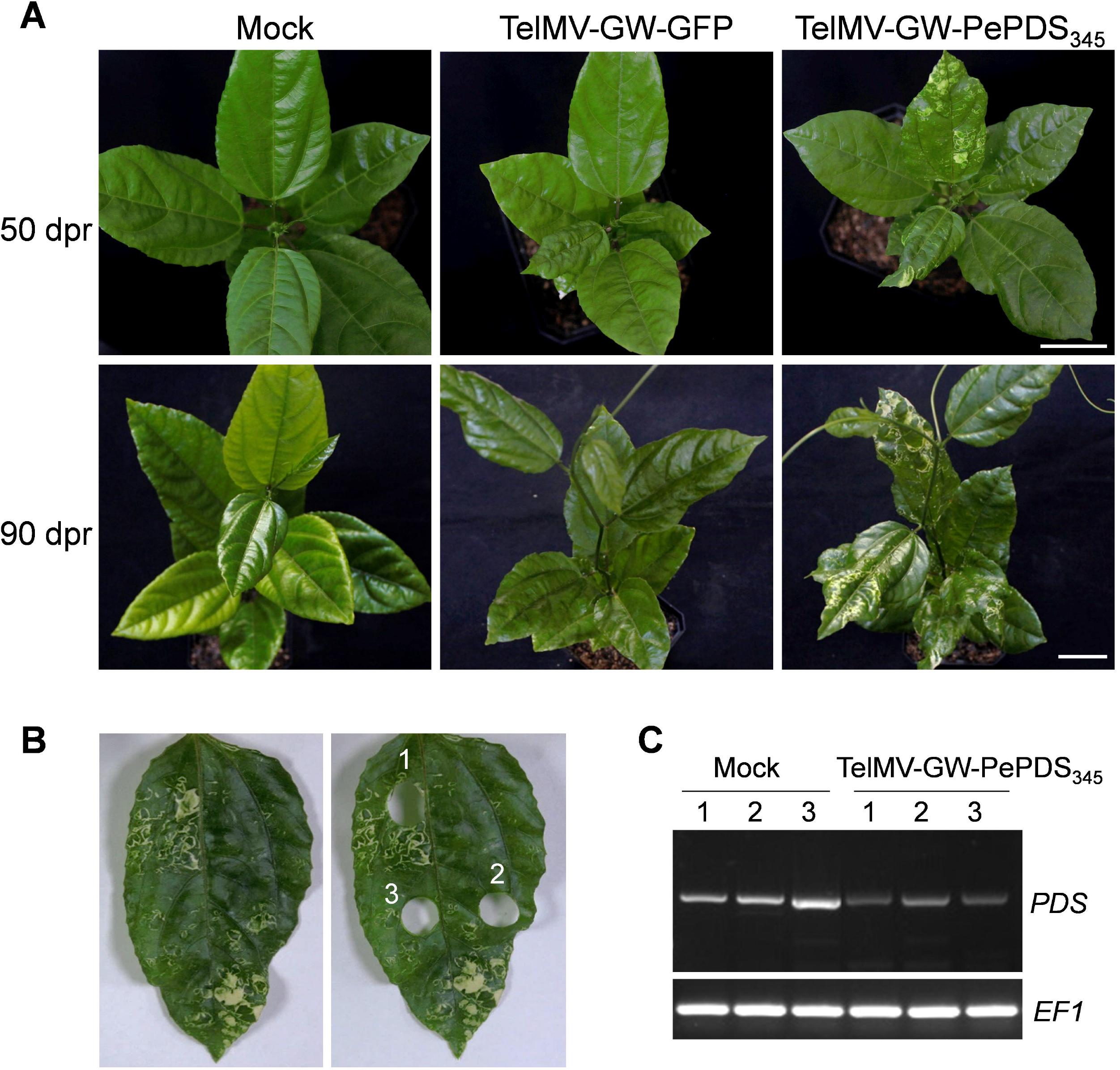
Endogenous gene silencing of *phytoene desaturase* (*PDS*) in passion fruit plants using TelMV-based VIGS vector. (A). Passion fruit plants were inoculated with buffer (Mock), TelMV-GW-GFP or TelMV-GW-PePDS_345_. Photos are taken at 50 and 90 days post rub-inoculation (dpr), respectively. Bars, 5 cm. (B). Schematic representation of passion fruit leaf inoculated by TelMV-GW-PePDS_345_ for sampling for RT-PCR analysis (left: before sampling; right: after sampling). (C). RT-PCR analysis of *PDS* mRNA level sampled from passion fruit leaf exhibiting bleaching phenotype. The passion fruit *EF1* gene serves as an internal control.

Next, RT-PCR was performed to quantify the *PDS* mRNA level using the *EF1* gene as the internal control. We selected three leaf spots (1, 2, and 3) representing the degree of photobleaching under normal night (Figure 4B). Total RNA of passion fruit showing bleaching on different spots was extracted and the subsequent RT-PCR analysis showed that the *PDS* mRNA level in the bleached leaf was indeed significantly decreased, compared to the Mock (Figure 4C). These observations suggest that TelMV-GW-PDS_345_ successfully triggered the endogenous gene silencing of *PePDS* in passion fruit plants.

## Discussion

This study reports the development of a passion fruit-infecting telosma mosaic potyvirus (TelMV)-based VIGS system for endogenous passion fruit gene silencing. The TelMV-based VIGS tool was proved to successfully silenced *PePDS* in passion fruit plants, resulting in a photobleached phenotype. To our knowledge, this is the first report on VIGS vector in passion fruit plants, and it provides a valuable tool for gene function analysis in passionfruit plants.

Virus-induced gene silencing (VIGS) is a powerful tool for gene function studies in plants. However, there was no VIGS tool available for passion fruit plants before this study. This is possibly due to TRV infection failure in passion fruit. In our preliminary tryout, we tested whether the most commonly used TRV vector could be utilized for VIGS in passion fruit. Unfortunately, it showed that TRV could not infect passion fruit plants using the agroinfiltration or rub-inoculation method in our current experimental setup (data not shown). Another reason is that the passion fruit infecting-virus infectious clone/vector has been reported recently, including TelMV, Passiflora mottle virus (PaMoV) and East Asian passiflora virus (EAPV) (Gou et al., 2023; Do et al., 2023; Chong et al., 2023). We expected that these newly constructed passion fruit infecting-virus infectious clones would be utilized as viral tools for protein expression and gene silencing in passion fruit plants.

Despite that *Potyvirus* is the largest group of plant RNA viruses and comprises many agriculturally important viruses, few potyviruses have been reported to be used as VIGS vectors for plant endogenous gene silencing. This is mainly due to two reasons, one is that potyvirus encodes a strong RNA silencing suppressor HC-Pro that inhibits the host RNA-based antiviral response (Valli et al., 2018). With this regard, a mild passion fruit-infecting potyvirus strain harboring HC-Pro point mutation exhibiting weakened or abolished RNA silencing suppressor activity would contribute a lot towards efficient gene silencing in passion fruit plants using potyvirus-based VIGS vector. Secondly, potyvirus employ polyprotein processing as its gene expression strategy, pioneers in potyvirus have demonstrated the manipulation of the potyviral genome is restricted and the mainly used foreign fragment insertion sites were P1/HC-Pro and NIb/CP (Xie et al., 2021; Houhou et al., 2021).

To achieve the fast, simple, and efficient manipulation of the potyviral genome, herein we inserted the Gateway recombination sites into the potyviral NIb/CP intercistronic sites and subsequently demonstrated the successful expression of GFP in this potyviral vector (Figure 2), suggesting our TelMV-GW could be used as protein expression vector. It is well known that the ability to induce endogenous gene silencing depends on the ability of the VIGS vector to systemically infect the plant of interest. In Figure 2, we clearly show that TelMV-GW-GFP successfully induced systemic infection both in model plant tobacco and passion fruit. This result further supports the TelMV could be used as a VIGS tool for endogenous gene silencing in passion fruit plants. In our study, we observed that TelMV-GW-GFP as well as TelMV-GW-PDS_345_ does not compromise the growth of passion fruit plants and the plants did not display severe viral symptoms such as stunting (data not shown), leading us to speculate that TelMV-VIGS vector is a mild strain compared to wild-type TelMV. Further experiments including phenotype analysis and molecular detection such as qPCR and Western blot are needed to validate this assumption.

Furthermore, the TelMV-GW vector was revealed to be used as a gene silencing vector both in *N. benthamiana* and *P. edulis* plants. This is supported by 1) TelMV-GW containing GFP fragments triggered gene silencing in GFP-transgenic tobacco plants (16c) (Figure 3), and 2) TelMV-GW expressing a partial fragment of *phytoene desaturase* (*PDS*) gene resulted in photobleached phenotype correlated with the reduced *PDS* mRNA levels (Figure 4). In line with this, two cases of potyvirus-based VIGS vectors have been reported recently. In 2021, Shen laboratory reported the papaya leaf distortion mosaic virus (PLDMV)-based VIGS vector for silencing genes in papaya, they successfully silenced five endogenous papaya genes, including, *PDS, Mg-chelatase H subunit*, putative *GIBBERELLIN (GA)-INSENSITIVE DWARF1A and 1B* and the cytochrome P450 gene *CYP83B1* (Tuo et al., 2021). Another work from the Stewart laboratory reported the maize dwarf mosaic virus (MDMV)-based VIGS vector for silencing endogenous maize genes (Xie et al., 2021). In addition, a preprint from Daròs laboratory demonstrated a mild isolate of watermelon mosaic virus (WMV)-based VIGS in cucurbits (Houhou et al., 2021). Taken together, these reports illustrated that potyviruses can also be used for VIGS in multiple plant species, including papaya, maize, melon and passion fruit. Nevertheless, more experiments should be carried out to illustrate the capability of the TelMV-based VIGS vector for silencing various passion fruit endogenous genes, including common VIGS marker genes, such as *Magnesium chelatase subunit I* (*CHLI*) and *Mg-chelatase H subunit*.

## Materials and Methods

### Plant materials and growth conditions

*N. benthamiana* or *P. edulis* Sim seeds were sown in 10-cm-wide square pots at two-cm depth and germinated in PINDSTRUP substrate supplemented with and vermiculite and perlite (4:1:1). Individual plants were transferred to pots and grown in a growth chamber with 16 h of light at 25°C and 8 h of darkness at 23°C. The relative humidity with set at 65% relative humidity.

### Plasmid construction

TelMV-GW: The plant expression binary vector TelMV-GW was based on the previously reported viral vector pPasFru (Gou et al., 2023). The Gateway recombination frame (Fragment 2, F2) was amplified from pEarleyGate 103 (Invitrogen, CA, USA) using the primer pair c/d (Table 1), and subsequently inserted between the NIb and CP cistron by the following overlapping PCR method: Fragment 1 and 3(F1, F3) were amplified from pPasFru using the primer pairs a/b and e/f (Table 1), respectively. F1 contains the viral sequence starting from the *Spe*I site on 6K2 to the end of NIb. F3 comprises the sequence from the end of NIb to the *Avr*II site on the CP cistron. Furthermore, F2 was inserted between the F1 and F3 by the overlapping PCR using primer pair a/f. The resulting fragment was named F4. Lastly, F4 was double digested with *Spe*I and *Avr*II enzymes and subsequently ligated into the predigested backbone vector pPasFu to construct the Gateway-based TelMV vector TelMV-GW.

**Table 1.**
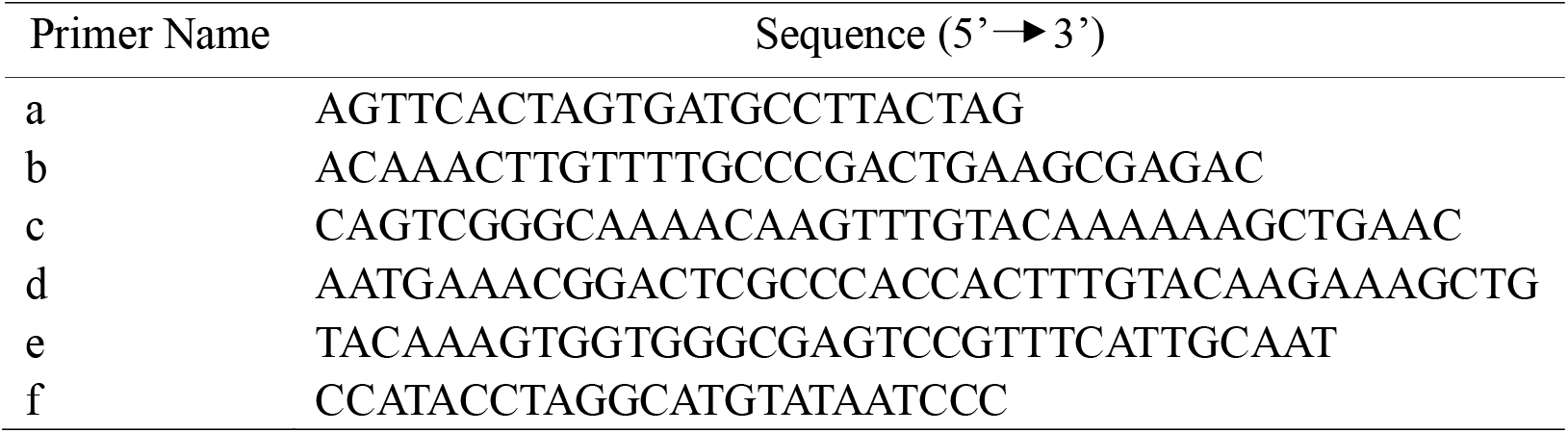
Primers used for TelMV-GW construction.

TelMV-GW-GFP: The full length of GFP gene was amplified from GFP-tagged TelMV infectious clone (TelMV-GFP) (Gou et al., 2023) using primer pairs GFP-F (5’ GGGGACAAGTTTGTACAAAAAAGCAGGCTTCATGAGTAAAGGAGAAGAA CTTTTC3’) and GFP-R (5’GGGGACCACTTTGTACAAGAAAGCTGGGTCGTCGTCCTTGAAGAAGATGGT3’). The PCR product was then cloned into pDONR221 (Invitrogen, CA, USA) through the BP reaction and subsequently cloned into the destination vector TelMV-GW by LR reaction.

TelMV-GW-GFP309 and 405: The full length of GFP was divided into two fragments GFP309 and GFP405 comprising 309 bp and the remaining 405 bp, respectively for amplification using TelMV-GFP as the template and primer pairs GFP309-F (5’ GGGGACAAGTTTGTACAAAAAAGCAGGCTTCATGAGTAAAGGAGAAGAA CTTTTC3’)/GFP309-R (5’GGGGACCACTTTGTACAAGAAAGCTGGGTCGTCGTCCTTGAAGAAGATGGT3’) and GFP405-F (5’GGGGACAAGTTTGTACAAAAAAGCAGGCTTCGGGAACTACAAGACACGTGCT3’) and GFP405-R (5’ GGGGACCACTTTGTACAAGAAAGCTGGGTCTTTGTATAGTTCATCCATGCCA TG 3’). The GFP309 and GFP405 fragments were recombined into pDONR201 (Invitrogen, CA, USA) using the BP CLONASE enzyme, followed by the LR reaction using TelMV-GW as the destination vector.

TelMV-GW-PDS_345_: As the passion fruit PDS gene (*PePDS*) has not been reported previously, we first did the mining in the genome sequence of *P. edulis* by BLAST searching using either tobacco PDS gene (*NbPDS*) or Arabidopsis PDS gene (*AtPDS*) as the query. Next, multiple sequence alignments were performed among the putative *PePDS, NbPDS* and *AtPDS*. A primer pair for amplifying *PePDS* fragment (345 bp) was designed from the conserved nucleotide sequence of *PePDS* and used for RT-PCR of *P. edulis* leaf. The PCR product was further used for cloning to the destination vector TelMV-GW through classic BP and LR reactions.

### Plant Inoculation

For the inoculation of *N. benthamiana* plants, *Agrobacterium tumefaciens* harboring virus infection clones were used for leaf infiltration. In brief, *A. tumefaciens* strain GV3101 harboring various plasmids was grown overnight on a shaker at 200 rpm and the bacterial culture was then centrifuged, washed and finally resuspended in the agroinfiltration buffer (10 mM MgCl2, 10 mM MES, pH 5.7) supplemented with 200 μM acetosyringone. The optical density (OD600) of the bacterial suspension was then adjusted to 0.5 for infiltration using a needle-less syringe.

For the inoculation of *P. edulis* plants, we used the rubbing method. More specifically, the sap prepared from *N. benthamiana* plants infiltrated with agrobacterium harboring relevant plasmid was rub-inoculated on the cotyledons of *P. edulis* plants.

### RT-PCR

Total RNA was extracted from 30 mg leaf tissue of *Nicotiana benthamiana* or *P. edulis* plants using TRNzol universal reagents (Tiangen). For first-strand cDNA synthesis, 500 ng of RNA was treated by DNase I (Thermo Fisher Scientific), followed by reverse transcription reaction using the SuperScript III First-strand Synthesis System (Thermo Fisher Scientific) as instructed. For RT-PCR, primers pairs TelMV-8285-F (5’ CCAAAGTTAGAGCCAGAAAGGA3’)/TelMV-8900-R (5’GAAACCATTCATGAC GACATTCA3’) and GFP-1-F (5’ATATAGGATCCATGGCAATGAGTAAAGGAGAAGAACTTTTCAC3’)/GFP-1-R (5’TTTGTATAGTTCATCCATGCCATG3’), were used to detect the viral mRNA and GFP mRNA, respectively. A fragment of *EF1* was amplified from passion fruit cDNA using primers EF1-F (5’GGCTGAGCGTGAACG TGGTA 3’)/EF1-R (5’CGGCACAATCAGCCTGGGAA3’) and served as the internal control. Primers Actin-F (5’AAAGACCAGCTCATCCGTGGAGAA3’) and Actin-R (5’ TGTGGTTTCATGAATGCCAGCAGC3’) were used for amplifying a fragment of *N. benthamiana Actin* gene.

## Data Availability Statement

The original contributions presented in the study are included in the article/Supplementary Material, further inquiries can be directed to the corresponding author/s.

## Author Contributions

ZD, HC, and XW designed the experiment and wrote the manuscript. XW performed the experiments. All authors analyzed, discussed the data, read, and approved the final manuscript.

## Acknowledgments

We thank Dr. Wenping Qiu (Missouri State University, USA) for revising the manuscript, Dr. Justice Norvienyeku (Hainan University, China) for helpful discussion and Dr. Runmao Lin (Hainan University, China) for helping the mining in the genome sequence of *P. edulis* for *PDS* gene.

